# Chemokine-biased robust self-organizing polarization of migrating cells *in vivo*

**DOI:** 10.1101/857623

**Authors:** Adan Olguin-Olguin, Anne Aalto, Benoît Maugis, Michal Reichman-Fried, Erez Raz

## Abstract

The mechanisms facilitating the establishment of front-rear polarity in migrating cells are not fully understood, in particular in the context of bleb-driven directional migration. To gain further insight into this issue we utilized the migration of zebrafish primordial germ cells (PGCs) as an *in vivo* model. We followed the molecular and morphological cascade that converts apolar cells into polarized bleb-forming motile cells and analyzed the cross dependency among the different cellular functions we identified. Our results underline the critical role of antagonistic interactions between the front and the rear, in particular the role of biophysical processes including formation of barriers and transport of specific proteins to the back of the cell. These interactions direct the formation of blebs to a specific part of the cell that is specified as the cell front. In this way, spontaneous cell polarization facilitates non-directional cell motility and when biased by chemokine signals leads to migration towards specific locations.

## Introduction

Establishing and maintaining polarity is essential for a range of different cellular functions^1^. This requirement is particularly critical for single-cell migration, a process that depends on the definition of a front-rear axis, which, in turn, dictates the direction of movement. This important issue has previously been investigated primarily in cells such as leukocytes, fibroblasts, *Dictyostelium* and neural crest cells^2–7^, cell types that employ actin polymerization to generate physical force to advance their leading edge. These studies suggest that polarity is generated and stabilized via the processes of local positive feedback control loops, long-range negative feedback loops and control protrusion formation by membrane tension, thereby allowing the cells to move forward^8–11^.

In contrast to actin-powered lamellipodia-based migration, a different cell migration strategy exhibited by diverse cell types in physiological contexts as well as in pathological conditions is characterized by the generation of bleb-type protrusions at the cell front^12–14^. Blebs are spherical in shape and are powered by hydrostatic pressure and cytoplasmic flow^15–17^. Here, a net forward movement is achieved via polarized bleb formation driven by inflow of cytoplasm and concomitant retraction at the opposite side of the cell. Several processes and molecular activities have been found to contribute to the generation of blebs. Most notably, actomyosin contractility is required to generate the intracellular hydrostatic pressure and to cause the breaks in the cell cortex that together allow the plasma membrane to separate from the underlying actin filaments^18^. Nevertheless, little is known about the mechanisms responsible for defining the front of blebbing cells and establishing the rear. Similarly, the nature of the interactions between the front and the rear are not well understood in this context.

An important model for studying bleb-driven motility in an *in vivo* context is that of zebrafish primordial germ cells (PGCs)^19^. Zebrafish PGCs migrate within the embryo using blebs, and their motility is normally directed by the chemokine Cxcl12a and its receptor Cxcr4b^20^. Whether directed by the chemokine or not, during their migration, PGCs alternate between two distinct modes of behavior namely ‘run’ and ‘tumble’^21^. During ‘run’ phases, PGCs extend blebs in the direction of movement and actively migrate, while during the ‘tumbling’ phases, PGCs are morphologically apolar and form blebs in all directions^21^. This periodic sequence of events where PGCs lose and regain polarity makes them an excellent model for studying the establishment and maintenance of cell polarity of blebbing cells and for determining the role of chemokine signaling in this process.

## Results

### Polarized distribution of molecules and structures along the front-rear axis of migrating PGCs

To shed light on the events and cross-interactions among cellular elements that facilitate cell polarization, we first identified proteins and structures that are asymmetrically distributed in migrating PGCs. The front of migrating PGCs exhibit blebs and contain a dense F-actin structure (termed actin brushes) (Fig. 1a, see also ^22^). Detailed analysis of the distribution of blebs around the cell perimeter revealed that they preferentially form in the direction of migration, with less frequent formation of smaller blebs at the cell rear (Supplementary Fig. 1). Next, with the aim of identifying additional elements that are distributed unevenly along the front-rear axis of the polarized cell, we set out to determine the position of organelles and molecules. Interestingly, while the endoplasmic reticulum (ER) is found throughout the cell body, it is absent from expanding blebs at the cell front (Fig. 1b). Two other structures, the microtubule organising center (MTOC, visualized using a microtubules (MTs) marker) and the Golgi apparatus (Fig. 1c and 1d, respectively) reside at the rear of the cell, behind the nucleus. Importantly, Ezrin, a member of the ERM family that links the plasma membrane to the actin cortex, is strongly enriched at the rear of the cell (Fig. 1e), similar to findings in some other cell types^23,24^. Interestingly, extended synaptotagmin 2a (Esyt2a), which belongs to a family of proteins linking the ER to the plasma membrane^25,26^ is also localized to the rear of the cell (Fig. 1f). Last, we found that Septin9a is enriched at the cell rear (Fig. 1g), where it might act in controlling cortical rigidity (reviewed in ^27^). Taken together, we detected actin accumulation at the cell front where blebs form, and, importantly, we observed localization of molecules that could potentially limit the formation of blebs at the rear of the cell.

**Fig. 1.**
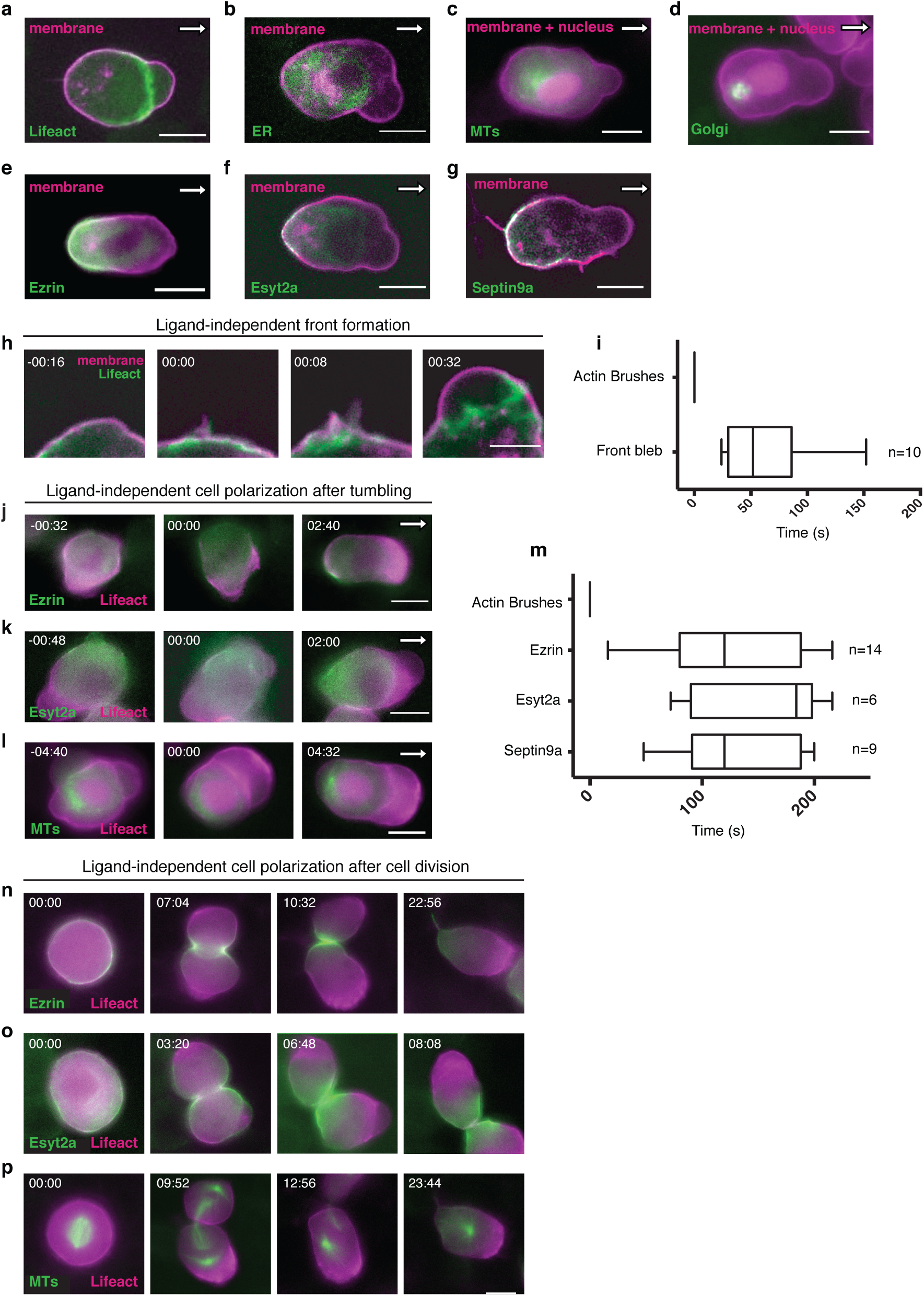
Polarized distribution of proteins and subcellular structures within migrating primordial germ cells (PGCs) and steps of self-organized ligand-independent polarity. **a-g** the localization fluorescent protein fusions of **a** Lifeact, **b** ER (Calreticulin signal peptide), **c** Microtubules (MTs, Clip-170H), **d** Golgi (N-terminus of Beta 1,4- galactosyltransferase), **e** Ezrin, **f** Extended synaptotagmin-like 2a (Esyt2a) and **g** Septin9a, relative to the plasma membrane (Farnesylation signal from c-Ha-Ras fused to fluorescent protein (magenta)), as well as the nucleus (**c** and **d** (magenta)) in wild type cells. White arrow indicates the direction of migration. Scale bars 10 *µ*m. **h** The formation of actin brushes at the new front and inflation of the first bleb. Times in minutes and seconds. Scale bar 5 *µ*m. **i** The time of bleb formation relative to the establishment of actin brushes at the cell front (median, 5 and 95 percentiles are indicated. n= number of cells. time in seconds). **j-l**, The establishment of the cell rear in the absence of receptor signaling as judged by the distribution of **j** Ezrin, **k** Esyt2a and **l** MTs (Clip170H) in green. The forming cell front is marked by Lifeact (magenta). Left panels in **j-l** present apolar tumbling cells, middle panels show the appearance of Actin within the new cell front (time 00:00) and in the right panels the localization of the proteins at the rear of the cell is presented. Times indicate minutes and seconds before and after the establishment of the front and the white arrows point at the direction of migration. Scale bar 10*µ*m. **m.** Time of ligand-independent polarization of different rear markers relative to the time of actin localization at the cell front (median, 5 and 95 percentiles are presented, n= number of cells, time in seconds). **n-p** Polarity establishment following cell division. Lifeact presented in magenta and the different rear markers in green (**n** Ezrin, **o** Esyt2a and **p** MTs (Clip170H). Left panels show the apolar cells prior to cytokinesis and subsequent panels present the progression through cell division, with the right panels showing the polarized daughter cells. Scale bar 10*µ*m.

### Definition of the cell front precedes the formation of the rear during exit from tumbling in ligand-independent PGC polarization

As a first step in exploring the mechanisms responsible for the establishment of polarity in the migrating cells, we determined the temporal course of events by which apolar tumbling cells establish the distribution of polarity markers. This analysis was first performed under conditions where the guidance cue Cxcl12a (the ligand of the chemokine receptor Cxcr4b) was absent (hereafter referred to as “ligand-independent cell polarization”) and then under conditions where the cells were presented with a polarized chemokine source.

Interestingly, whereas blebs are dramatically different from actin-powered lamellipodia regarding their molecular composition and morphology, we found that similar to findings in mesenchymal cells^28^, actin polymerization is the earliest detectable marker of the future front of PGCs (considered “time 0”, with the first bleb forming, an average of 62 seconds after accumulation of actin in the front (Fig. 1h, I, median= 52 seconds). Importantly, based on the localization of Ezrin, Esyt2a, and Septin9a, we found that markers of the cell’s rear developed more than 120 seconds after the appearance of the actin brushes (Fig. 1j, k, m, Supplementary Fig. 2 and Supplementary Movie 1). This observation is consistent with the idea that the position of the cell rear is dictated by the location at which the front forms. The positioning of the MTOC at the cell rear does not always occur within the period we followed and represents a late event in PGC polarization. Thus, the MTOC is unlikely to be involved in the initial development of the front-rear axis (Fig. 1l).

### The rear of the cell is defined prior to the cell front following cell division in ligand-independent PGC polarization

Another key stage at which PGCs become polarized is at the end of cell division. During mitosis, the cells assume a round morphology and as judged by the distribution of the different molecules, the cells are apolar (Fig. 1n-p, time point 00:00). Interestingly, the cleavage furrow that develops during the last stages of mitosis harbors proteins we found to reside at the cell rear. Accordingly, Ezrin and Esyt2a are the first proteins to become localized along the front-rear axis upon repolarization (Fig. 1n, o). In the absence of a chemoattractant, the cell front as marked by actin brushes formed away from the rear marked by Ezrin/Esyt2a (Fig. 1n,o, Supplementary Movie 2). In this scenario, the MTOC that is located initially at the cell front translocate to the cell back (Fig. 1p time point 23:44, Supplementary Movie 2). These results suggest that while the cell front initiates polarization when PGCs exit from tumbling, the rear of the cell can also direct cell polarization and thereby determine the location of the front, as seen following cell division.

### The establishment the front-rear axis in blebbing cells by localized Rac1 activity

Since we found that as PGCs exit from tumbling the front is established before the rear, we sought to determine the role of actin polymerization in the establishment of the front-rear axis. To this end, we first studied the dynamics of Rac1 activation in the absence of the chemoattractant. To follow active Rac1 protein distribution, we used the GTPase binding domain (GBD) of PAK-1 protein^29^ as the PGCs acquire polarity. Interestingly, we observed the formation of a crescent of Rac1 activation (Fig. 2a, time point - 00:16) preceding the formation of the actin brushes (Fig. 2a, time point 00:00) and of other polarized molecules and structures. Rac1 activation followed by actin accumulation can then be observed at the membrane of newly formed blebs at the cell front, thereby maintaining the front-biased actin accumulation (Fig. 2a, time point 00:16).

**Fig. 2.**
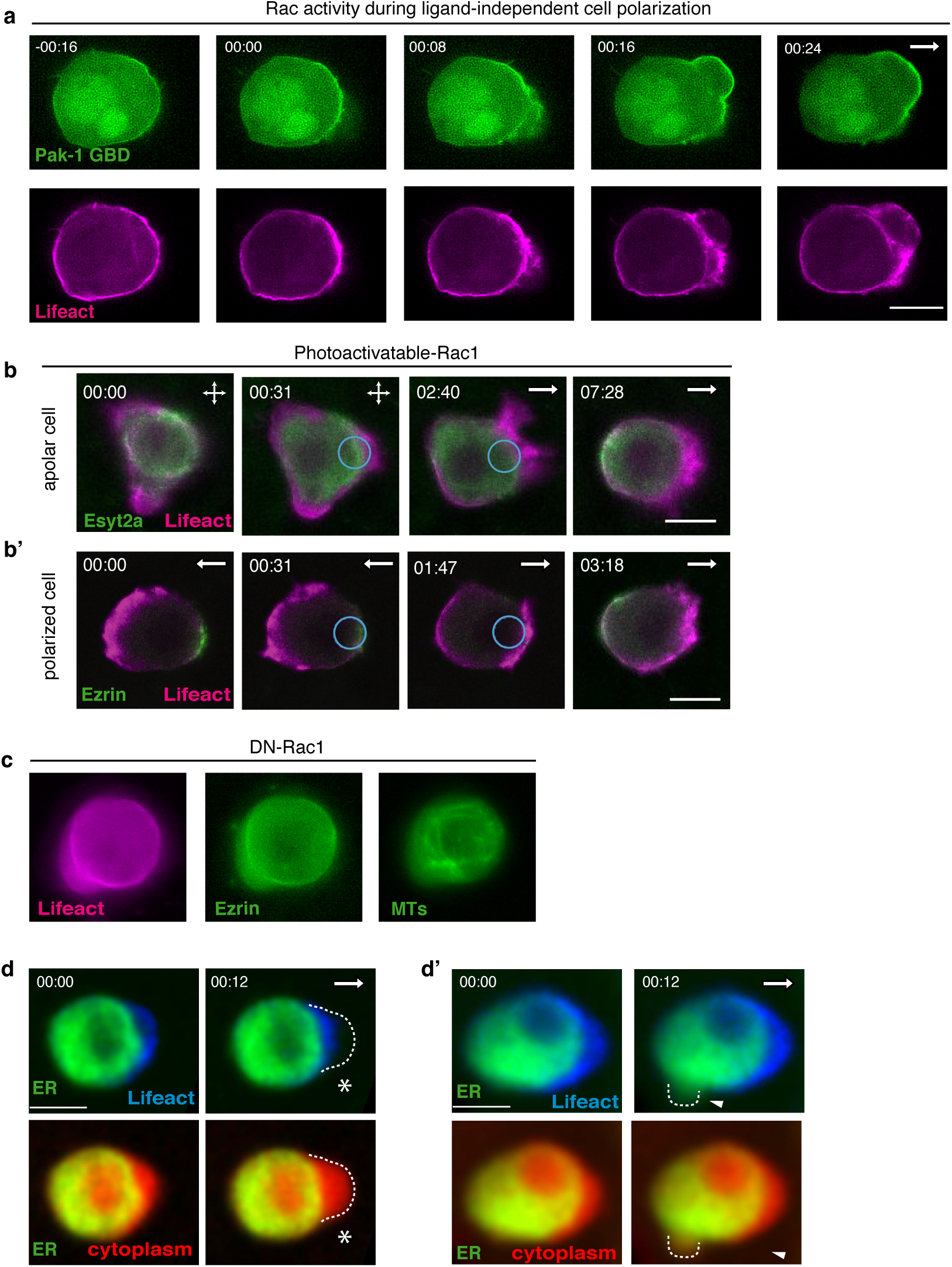
The role of actin brushes and Rac1 GTPase in the establishment of the self-organizing polarity in PGCs. **a** Polarization of GBD of PAK-1 fluorescent protein fusion (green), marking locations in the cell where Rac1 is active in the absence of receptor signaling. Actin brushes in the newly established front (magenta) are presented for the same time points. **b** Light mediated Rac1 activation is sufficient for inducing polarization in tumbling PGCs (**c**) and for reversing it in polarized cells (**b’**). Lifeact is presented in magenta and Esyt2a and Ezrin fusion proteins in green. Circles indicate the area where the light inducible Rac1 was activated. **c** Rac1 activity is required for polarity establishment as judged by lack of stable focused accumulation of Lifeact (left panel) Ezrin (middle panel), and MTs (Clip170H, right panel) in cells expressing the dominant-negative form of Rac1 (DN-Rac1). **d** and **d’** Wild type PGCs labeled with Lifeact (blue), ER (green) and cytoplasm (red). The blebs are marked by the dashed line. The asterisk indicates the front bleb and arrowhead points at a bleb formed at the cell rear. Scale bars 10*µ*m. Time in minutes and seconds and white arrows indicate the direction of migration.

To examine whether formation of the front plays an instructive role in the definition of the cell rear, we employed a photoactivatable version of Rac1^30^ (Fig. 2b, Supplementary Movie 3). Indeed, activation of Rac1 in tumbling cells directed the formation of the front to the site of activation (Fig. 2b). Remarkably, activating Rac1 at the rear of already-polarized cells resulted initially in the formation of two actin-rich areas at the same time (Fig. 2b’). Subsequently, the area that exhibited a stronger actin signal was further maintained, followed by the formation of the rear at the opposite side of the cell. Specifically, following the generation of the front, Ezrin was enriched at the new forming rear (Fig. 2b’). These results thus show that the formation of the cell front is sufficient for positioning of the rear.

To determine the role of Rac1 in the polarization of PGCs, we inhibited its activity by expressing a dominant negative (DN) form of the protein (Rac1T17N)^31^, thereby abolishing the formation of the actin brushes at the cell front. As we have previously shown, under these conditions cells are not elongated, do not migrate, and exhibit apolar blebbing activity (Fig. 2c, Supplementary Movie 4, ^32^). This phenotype could result from weakening of the actomyosin cortex coupled with residual contractility, which leads to more frequent separation of the plasma membrane from the cortex and bleb formation. Importantly, under these conditions none of the molecular markers for the rear of the cell (Ezrin, MTOC) were polarized (Fig. 2c, Supplementary Movie 4). These results thus show that Rac activity is essential for the generation of the cell front and, under these conditions, it is also needed for the establishment of the rear.

In addition to the role of Rac1 activation in increasing blebbing activity, we found the formation of the actin brushes plays an additional role in the polarization of the cell. Monitoring the dynamic localization of different structures within the cells, we realized that the ER is excluded from blebs formed at the cell front (Fig. 1b, Fig. 2d). As actin brushes are found ahead of the ER, we speculated that they constitute a barrier blocking the entry of ER into the forming bleb (Fig. 2d). Consistent with this idea, the ER was capable of entering the blebs at the cell rear, a region devoid of actin brushes (Fig. 2d’). The smaller size of the blebs at the back of the cell (Supplementary Fig. 1b) could be, in part, a result of the more viscous ER^33^ that resists inflation by the cytoplasm, a scenario that is avoided at the cell front by the presence actin brushes.

### Actomyosin-based contractility is essential for the establishment of the cell of the rear

The finding that the formation of the actin brushes at the cell front was followed by the accumulation of the membrane-cortex and membrane-ER linker proteins and Septin9a at the cell back can potentially be explained by front-to-rear actomyosin-dependent retrograde flow^32,34–36^. To examine whether such a flow could be responsible for the formation of the cell rear in ligand-independent cell polarization, we expressed a dominant negative form of the Rho kinase protein (ROCK), which is known to inhibit actomyosin contraction (DN-ROCK)^22,37^. PGCs expressing the DN-ROCK were round and immotile and showed no polarization of the back markers (Ezrin, Esyt2a and MTOC) (Fig. 3a, Supplementary Movie 5). Strikingly, as judged by the Lifeact signal, the actin brushes characteristic of the cell front did form at one side of the cell. As we increased the contractility level, we observed an inverse phenotype; Expression of a constitutively active (CA) form of RhoA (RhoAG14V)^38,39^ resulted in strong retrograde flow and formation of a very large stable protrusion at the cell front (Fig. 3b, in agreement with findings in another cell type^38^. Consistent with the idea that the proteins for the rear of the cell are transported there via retrograde flow, we found strong localization of Ezrin, Esyt2a and MTOC at the rear of cells expressing the CA-RhoA protein (Fig. 3b and Supplementary Movie 6).

**Fig. 3.**
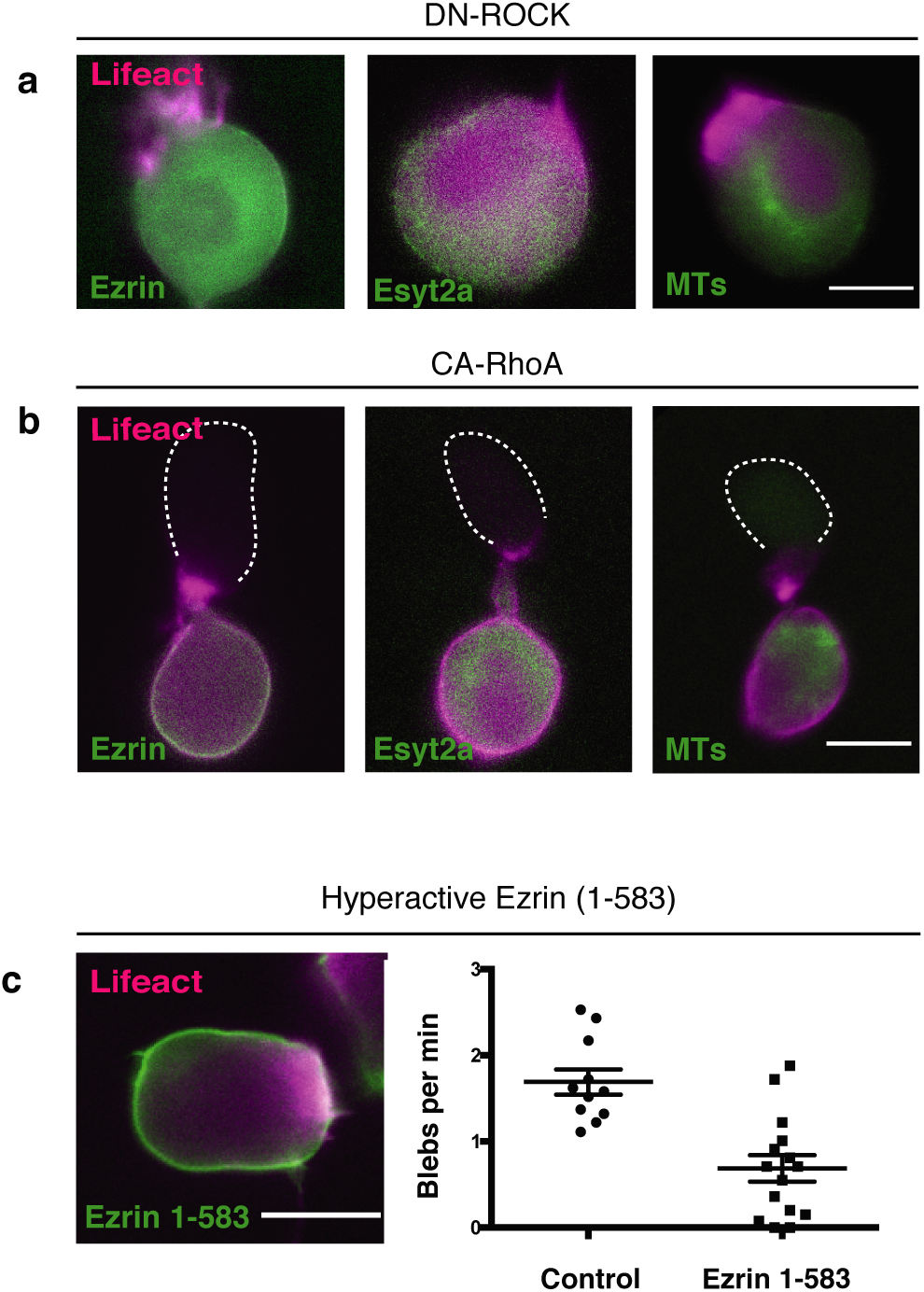
The role of contractility and of Ezrin function in the self-organized polarization of the PGCs. **a** ROCK activity is required for the establishment of the rear. Lifeact in magenta and Ezrin (left), Esyt2a (middle) and MTs (Clip170H, right) in green in cells expressing a dominant negative version of ROCK (DN-ROCK). **b** Enhanced contractility induced by expression of a constitutively active form of RhoA (CA-RhoA) results in hyper-polarization of the cell. Lifeact in magenta and Ezrin (left), Esyt2a, (middle) and MTs (Clip170H, right) in green. Dashed line markes the contours of the cell front. **c** Hyperactive version of Ezrin (amino acids1-583) is distributed over a larger portion of the cell perimeter (left panel, Lifeact in magenta and Ezrin in green) and inhibits blebbing activity (graph on the right, Ezrin (1-583) n = 15, Control n = 11, two tailed t-test with Welch’s correction, p= <0.0001). Scale bars 10*µ*m.

Next we sought to elucidate the functional relevance of Ezrin localization to the rear of the cell, and we specifically tested the possible role of the protein in preventing blebbing at the cell rear. Consistent with the idea that Ezrin inhibits bleb formation at the cell back, expression of a hyperactive version of Ezrin (Ezrin 1-583,^40^) strongly reduced blebbing activity around the cell perimeter (Fig. 3c). These findings show that an intermediate level of contractility is compatible with cell polarization and they show that Ezrin helps to define and stabilize the cell rear by preventing blebbing at this part of the cell, thereby favoring the formation of blebs at the other side.

### Orientation of front-rear cell polarity by chemokine signaling

The observations mentioned above and the results of the manipulations reveal that in PGCs intracellular polarity is established through a self-organizing mechanism that is independent of a guidance cue. We then proceeded to determine the mechanism by which a guidance cue directs the front-rear axis with respect to the position of the migration target. To this end, we first introduced a chemokine source next to tumbling apolar PGCs (Fig. 4a) and examined the course of polarity establishment. Interestingly, the order of the polarization events in cells responding to the chemokine were not different from those observed in self-organizing ligand-independent polarization (Fig. 4b-e and Fig. 1m). Specifically, we found that the actin brushes formed first, followed by the establishment of the rear as manifested by the accumulation of Ezrin, Esyt2a and MTOC (Fig. 4b-e and Supplementary Movie 7). Similar to the observation for the ligand-independent polarization, the positioning of the MTOC to the rear of the cell represented a late and temporally, highly variable event (between 2.5 to 42 minutes, Fig. 3e). This suggests that MTOC localization might be important for persistent migration but not for the initial polarization. These observations are consistent with the idea that the chemokine dictates the positioning of the cell front by biasing an underlying ligand-independent self-organizing polarity cascade.

**Fig. 4.**
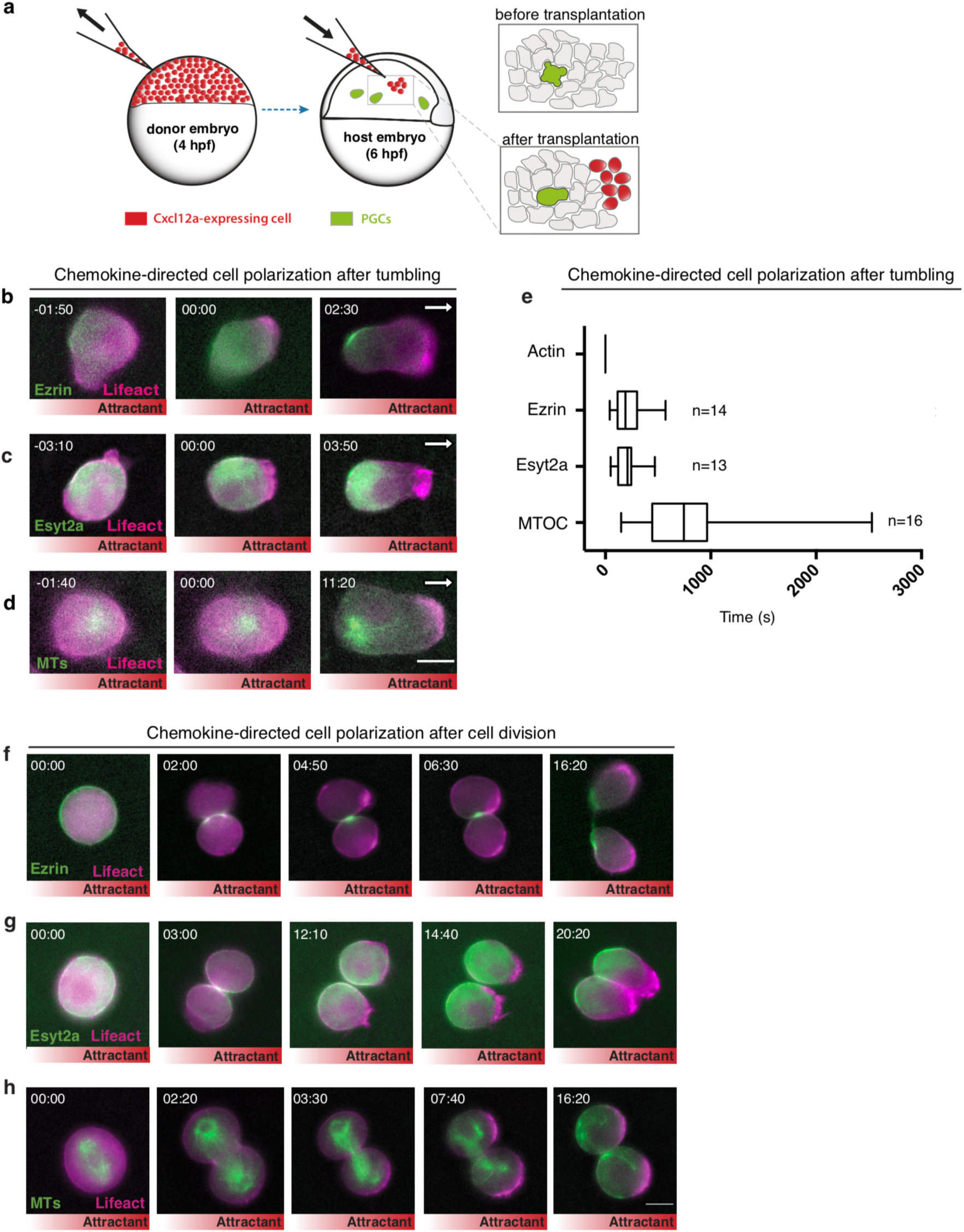
Chemokine-directed polarity establishment. **a** A scheme presenting the generation of local chemokine source within the live embryo. **b-d**, The forming cell front is labeled with Lifeact (magenta) and rear markers **b** Ezrin, **c** Esyt2a and **d** MTs (Clip170H) in green. Left panels show apolar cells, middle panels the appearance of the polarized front (time 00:00) and the right panels present the subsequent polarization of the rear markers. Time in minutes and seconds. White arrows indicate the direction of migration towards the chemokine source (red gradient below). **e** Timing of the chemokine induced polarization steps (median, 5 and 95 percentiles are presented, n= number of cells, time in seconds). **f-h**, Chemokine-directed cell polarization following cell division. Lifeact (magenta) and the rear markers **f** Ezrin, **g** Esyt2a and **h** MTs (Clip170H) in green. Left panels present the apolar cells prior to cytokinesis. In the second column one of the cells forms a front in the direction of the chemokine source, and in the third column the other cell forms a front as well. The right panels show the two polarized cells migrating in the direction of the chemokine source. In **b, c, d, f, g, and h** the distribution of the chemokine (Cxcl12a) is illustrated by the red gradient below the panels. Scale bars 10*µ*m.

Next, we examined the polarization of dividing PGCs while migrating in a chemokine gradient. According to our findings concerning the polarization of dividing cells in the absence of Cxcl12a, the rear formed at the cleavage plane thus determining the front at the opposite end. In the presence of chemokine gradient, while the rear markers first accumulated at the cleavage plane, the front formed at the side of the cell facing the higher level of the signal (Fig. 4f-h, Supplementary Movie 8). The position of the cell rear was then gradually adjusted according to the position of the new cell front.

To determine the mechanisms by which the activation of Cxcr4b, the receptor for Cxcl12a biases the self-organizing polarity cascade, we monitored the response of manipulated cells to a source of the guidance cue. Consistent with the idea that Cxcr4b activation leads to an increase in actin polymerization by controlling the Rac1 activation level^41,42^, germ cells in which Rac1 activity was inhibited did not polarize at the morphological level or at the molecular level in response to the chemokine (Fig. 5a, Supplementary Movie 9). Additionally, when contractility was inhibited by DN-ROCK, cells positioned within a chemokine gradient did not form a cell rear as judged by the distribution Ezrin and Esyt2a and the positioning of the MTOC (Fig. 5b, Supplementary Fig. 3). Intriguingly however, the majority of cells in which contractility was inhibited (39 out of 49) formed a clear actin-containing front in the direction of the source of the chemoattractant, indicating that they could sense the chemoattractant (Supplementary Fig. 3, Supplementary Video 10). These results show that receptor activation instructs the generation of the cell front, but in order to define a stable front-rear axis with a clear cell back, myosin-dependent retrograde flow has to localize proteins such as Ezrin and Esyt2a. As was the case in the absence of the ligand, increasing the contractility by expressing a constitutively active form of the RhoA protein (RhoAG14V, CA-RhoA) created a stable front bleb and pushed all the back markers and organelles to the rear (Fig. 5c, Supplementary Movie 11). In this case, however, the PGCs did not respond to the chemokine by directional migration towards the source (Fig. 5c’), which reflects the dominance of the strong retrograde flow-induced polarity and depletion of the receptor from the front bleb (Fig. 5d, d’).

**Fig. 5.**
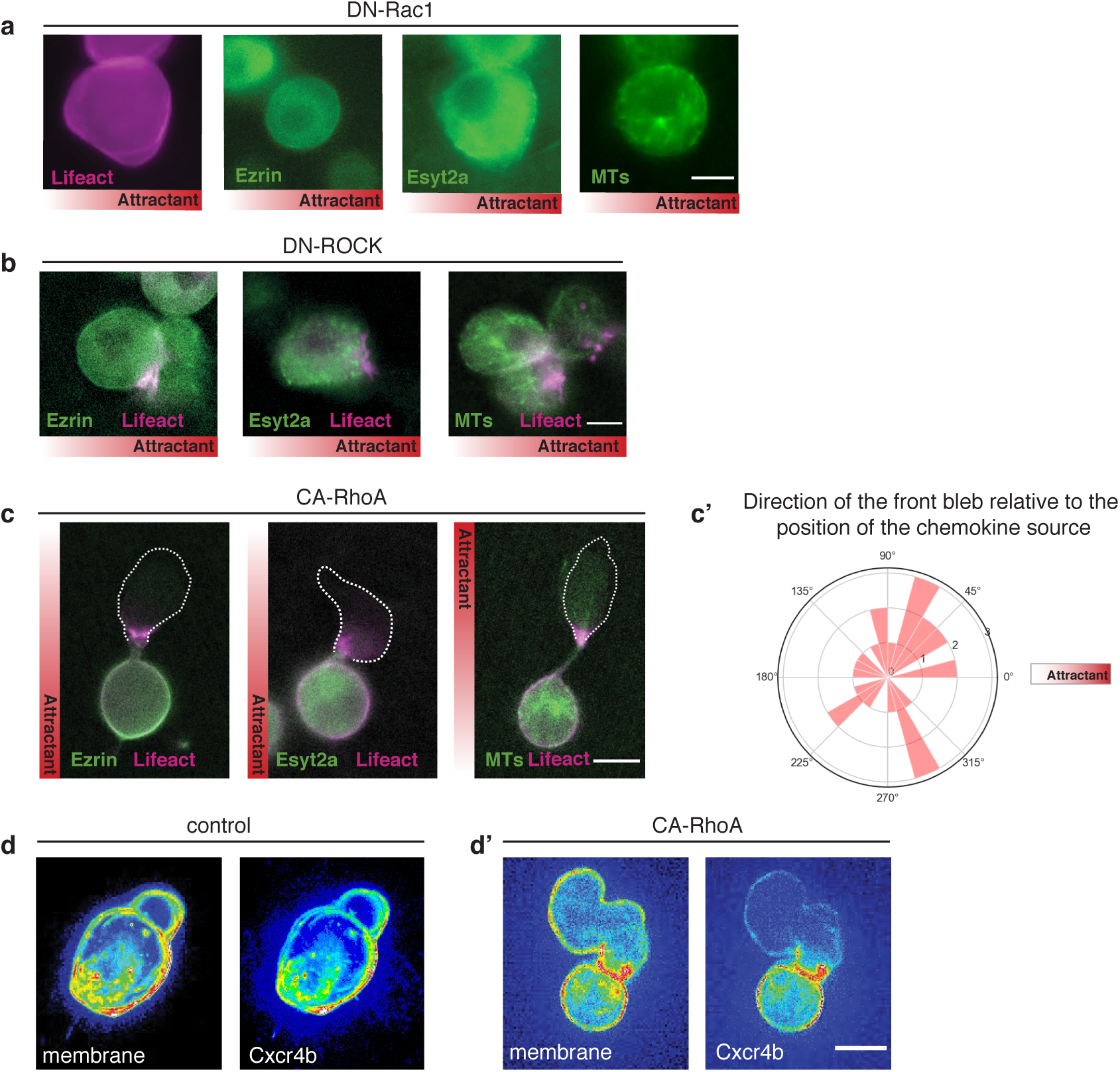
Mechanism of chemokine-induced polarization. **a** Rac1 activity is required for ligand-induced polarity establishment as judged by lack of stable focused accumulation of Lifeact (magenta), Ezrin (green, 2nd panel), Esyt2a (green, 3rd panel) and MTs (green, right panel) in cells expressing the dominant-negative form of Rac1 (DN-Rac1). **b** Inhibition of contractility by expression of a dominant negative version of ROCK (DN-ROCK) affects the definition of the cell rear. Lifeact in magenta and Ezrin (left panel), Esyt2a (middle panel) and MTs (Clip170H, right panel) in green. **c**, Enhancement of contractility by expression of a constitutively active version of RhoA (CA-RhoA) in the presence of the chemokine, Lifeact in magenta and Ezrin (left panel), Esyt2a (middle panel) and MTs (Clip170H, right panel) in green. **c’**, PGCs with increased contractility do not orient their protrusions towards the chemokine. Rose-plot, 0° indicates the source of the chemokine (n=25 cells). **d** In control cells Cxcr4b is expressed around the cell perimeter marked by the plasma membrane (z-projection). **d’** An increase in contractility decreases the amount of the receptor in the front protrusion (z-projection). In **a, b** and **c** the distribution of the chemokine (Cxcl12a) is illustrated by the gradient in the red rectangle. Scale bars 10*µ*m.

## Discussion

The temporal order and the mechanisms by which the front and rear of migrating cells are defined are not well understood, in particular in the context of the live tissue. In this work, we studied this issue in of bleb-driven migration. We identified polarized structures, organelles and molecules such as “linker proteins” (Ezrin, Esyt2a and Septin9a) that become localized to the rear of the cell and a polarized localization of the ER (Fig. 1). The microtubule-organizing center and the Golgi apparatus reside behind the nucleus, which differs from the from the position of these organelles in several other cell types ^43^.

Many of the studies on the polarization of migrating cells highlight the role of biochemical signaling pathways (e.g. small GTPases, GEFs, PI3P signaling)^41,42,44^ that define the front or the rear and the inhibitory interaction between them^8,45^. Our work underscores the importance of biophysical parameters in polarity establishment and maintenance in migrating cells. We report for the first time that actin polymerization at the cell front generates a barrier preventing organelles from entering the blebs, thereby controlling the composition of the expanding protrusion. By exclusion of organelles such as the ER, the inflation of the bleb could be less restricted, presumably contributing to the fact that blebs at the cell front are larger as compared with blebs formed in other locations of the cell (Supplementary Fig. 1). Another possible mechanism contributing to the polarization in bleb size along the front-rear axis could be the localization of the linker proteins Ezrin and of a molecule previously not known to reside at the cell back, Extended synaptotagmin-like protein 2a. These linker proteins, could play a role in polarizing bleb formation in migrating cells by connecting the plasma membrane to the ER and the actin cortex. While the *in vivo* model of PGC migration does not allow measuring membrane tension, this cellular feature has been shown to play a role in polarization of migrating cells^8,46^. Interestingly, this physical parameter can be regulated by Ezrin, which can enhance membrane tension^47^. Therefore, beyond preventing blebbing at the rear, Ezrin could additionally enhance cell polarity via control of membrane tension.

Our data reveals that during the establishment of cell polarity the front is defined first, with the rear forming at the other side of the cell a few minutes later in a process that depends on actomyosin contractility. Consistent with the reciprocal inhibition between the front and the rear of the cell, when the rear forms first, as is the case following cell division, the front is defined at the opposite side of the cell during self-organizing polarity establishment. The results we present are consistent with the idea that the role of the graded distribution of the chemokine is only to bias the initial definition of actin polymerization site that is followed by the development of the rear.

The importance of actin polymerization as a key determinant in polarity establishment in bleb-based cell migration is highlighted by the experiments in which it is inhibited by the expression of the dominant negative form of Rac1 and the activation of Rac1 activity at a specific part of the cell, which inhibits or is sufficient for inducing front-rear polarity respectively.

Considering the findings presented above, we propose the following model for self-organizing and chemokine-biased polarity establishment in bleb-driven single cell migration (Fig. 6a): The first step in the polarization cascade is the formation of actin brushes, which can be established at any point around the cell perimeter (6a, ii). In the absence of chemokine signaling, this position is determined stochastically. According to our model, the only role of the chemokine is to bias the formation of the actin brushes, such that they are more likely form in the part of the cell facing the higher level of the attractant source. The resulting increase in actomyosin contractility at the site where brushes form promotes breaks at the cortex, which defines weak points of membrane-cortex interaction and favors bleb formation at this aspect of the cell (6a, iii). The self-organizing polarization cascade then defines the rear of the cell by transport of linker proteins to the opposite side of the cell, where blebbing is inhibited. The result of this cascade is stable and robust polarization of the cell (Fig. 6b). This course of events raises a problem concerning *in vivo* cell migration as the robustness of the polarity generated precludes, or makes the introduction of corrections to the migration path very difficult. While the mechanistic basis for this phenomenon is currently unknown, changes in migration direction are made possible following “tumbling” phases, which are periodic events of loss of cell polarity^21^.

**Fig. 6.**
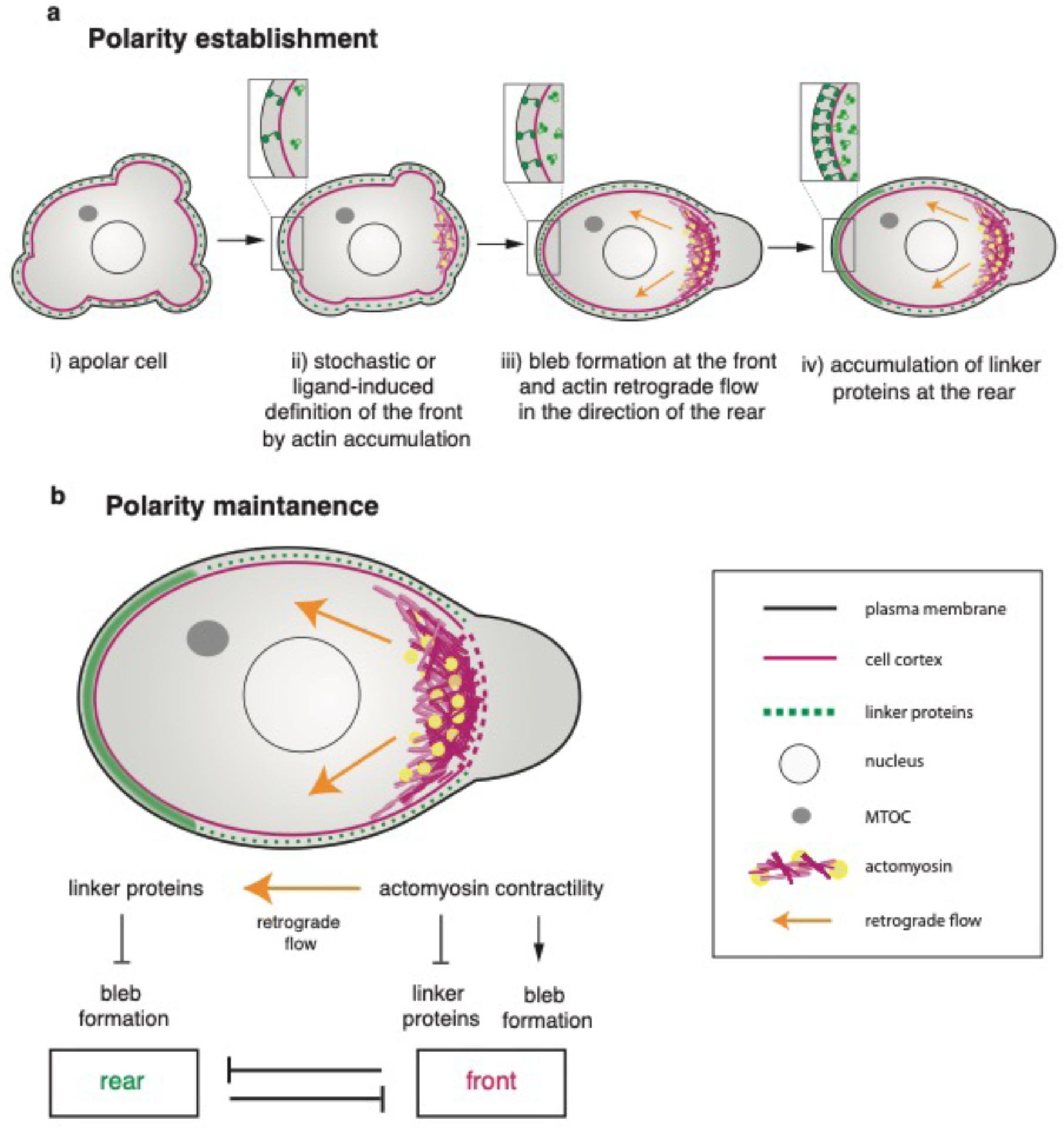
A model for polarity establishment and maintenance in migrating PGCs. **a** The exit from the apolar state (i) can either be self-organized, or induced by the polar distribution of the guidance cue. As judged by actin polymerization, the cell front is defined first (ii). Actin accumulation at the front focuses the protrusions to the same location and actomyosin dependent retrograde flow leading to accumulation of the linker proteins (Ezrin and Esyt2a, green) and the Septin9a protein at the rear (iii). High levels of linker proteins then accumulate at the cell rear (iv). **b** Polarity is maintained by the enhanced contractility at the cell front that promotes bleb formation at the front and flow of bleb-inhibiting linker proteins to the rear, where they inhibit bleb formation.

## Methods

### Zebrafish strains and fish maintenance

Zebrafish (*Danio rerio*) embryos of the AB and ABTL backgrounds and embryos from different transgenic fish lines (see Table 1) were used as wildtype embryos. Zebrafish were maintained according to the recommendations for the care of zebrafish in Annex A of the European Convention on the protection of vertebrates used for scientific purposes, EU Commission Recommendation 2007/526/EC and Article 33 of EU Directive 2010/63^48^.

**Table 1.**

| **Table 1** |  |  |
| --- | --- | --- |
| **Zebrafish strain used** | **transgenic ID** | **Reference** |
| wildtype strain of AB |  |  |
| wildtype strain of ABTL |  |  |
| *tg(kop:lifeact.EGFP.nos3'UTR-cry:DsRed)* | 38 | Hartwig et al, 2014 |
| *tg(kop:mCherry.F.nos3'UTR-cmlc:EGFP)* | 57 | Tarbashevich et al., 2015 |
| *tg(kop:GFP.F.nos3'UTR-cry:DsRed)* | 24 | Blaser et al., 2005 |
| *tg(kop:YPET.Ezrin.nos3'UTR-cmlc:EGFP)* | 99 | this work |
| *tg(kop:mCherry.Extended Synaptotagmin2al.nos3'UTR-cmlc:EGFP)* | 100 | this work |
| *tg(kop:mCherry.Clip170H.nos3'UTR-cmlc:EGFP)* | 101 | this work |
| *tg(kop:mTFP1.Septin9a.nos3'UTR-cmlc:EGFP)* | 106 | this work |
| *tg(kop:lifeact.mCherry.nos3'UTR-cry:DsRed)* | 118 | this work |
| *tg(kop:mCherry.F'.nos3'UTR-cmlc:EGFP; kop:lifeact.EGFP.nos3'UTR-cry:DsRed)* | 38/57 | this work |
| *tg(kop:GFP.F.nos3'UTR-cry:dsRed; kop:mCherry.Esyt2al.nos3'UTR-cmlc:EGFP)* | 24/100 | this work |
| *tg(kop:GFP.F.nos3'UTR-cry:dsRed; kop:mCherry.Clip170H.nos3'UTR-cmlc:EGFP)* | 24/101 | this work |
| *tg(kop:lifeact.EGFP.nos3'UTR-cry:DsRed; kop:mCherry.Esyt2al.nos3'UTR-cmlc:EGFP)* | 38/100 | this work |
| *tg(kop:lifeact.EGFP.nos3'UTR-cry:DsRed; kop:mCherry.Clip170H.nos3'UTR-cmlc:EGFP)* | 38/101 | this work |
| *tg(kop:YPET.Ezrin.nos3'UTR-cmlc:EGFP; kop:lifeact.mCherry.nos3'UTR-cry:DsRed)* | 99/118 | this work |
| *tg(kop:mTFP1.Septin9a.nos3'UTR-cmlc:EGFP; kop:lifeact.mCherry.nos3'UTR-cry:DsRed)* | 106/118 | this work |

### Generation of transgenic fish lines

Transgenic fish lines expressing either Ezrin, Esyt2a, Clip170H or Septin9a in PGCs, alone or in combination with Lifeact or a farnesylated fluorescent protein were generated for this study (see Table 2). Plasmids containing the kop promoter, the protein of interest (Lifeact, Ezrin, Esyt2a, Clip170H or Septin9a), with a fluorophore, the *nanos3* 3’UTR and a dominant marker were generated using Gateway Technology from Invitrogen. Purified plasmid was injected with RNA coding for Tol2.3 transposase into the cell of one-cell stage fish embryos and screened for the dominant marker. Fish carrying the *kop:lifeact.mCherry.nos3’UTR-cry:DsRed* were crossed to fish carrying the *kop:YPET.Ezrin.nos3’UTR-cmlc:GFP* and the fish carrying *kop:mTFP1.Septin9a.nos3’UTR-cmlc:EGFP* to generate the fish lines with both transgenes.

**Table 2.**
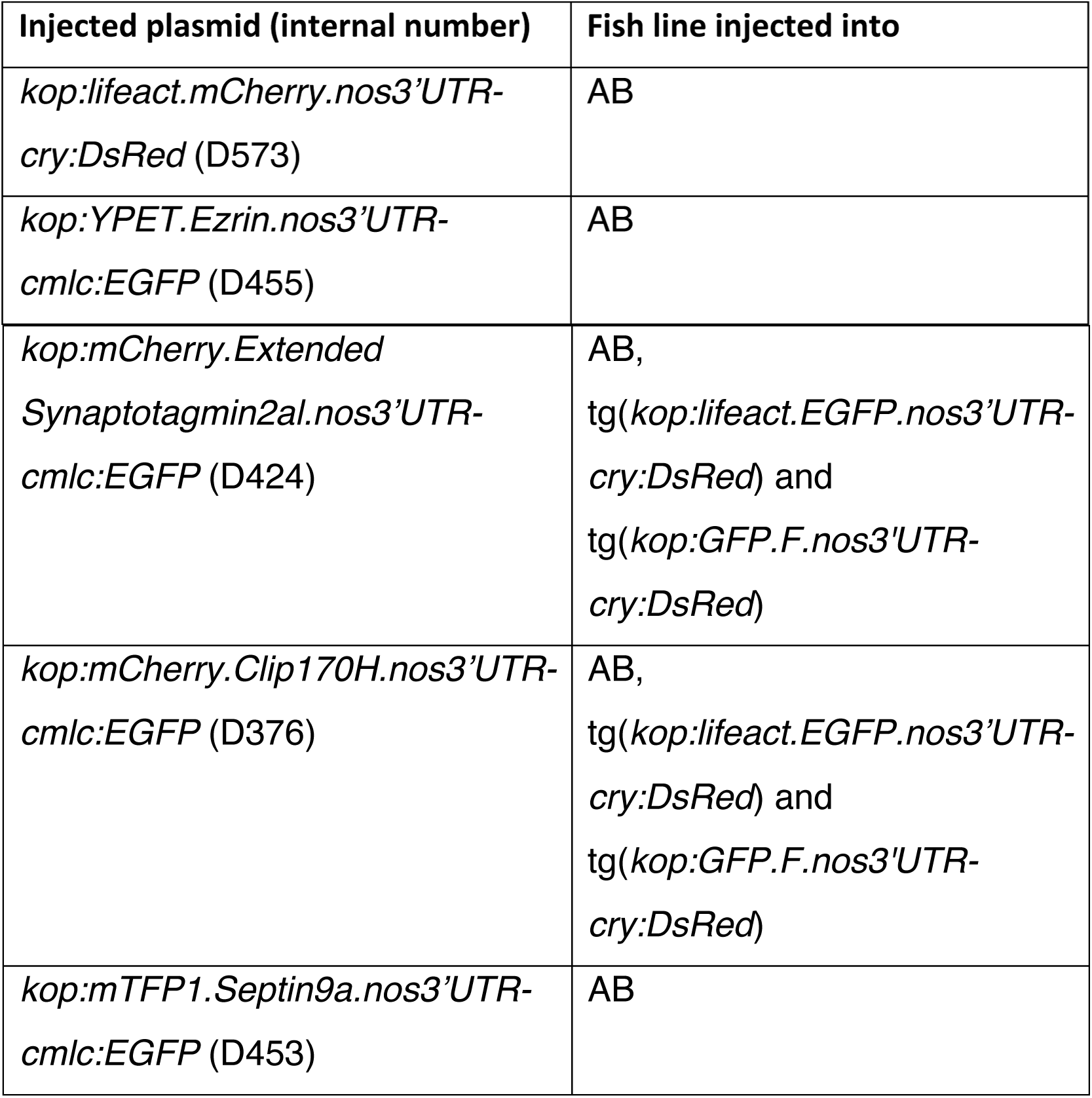

| Injected plasmid (internal number) | Fish line injected into |
| --- | --- |
| *kop:lifeact.mCherry.nos3'UTR-cry:DsRed* (D573) | AB |
| *kop:YPET.Ezrin.nos3'UTR-cmlc:EGFP* (D455) | AB |
| *kop:mCherry.Extended*<br>*Synaptotagmin2al.nos3'UTR-*<br>*cmlc:EGFP* (D424) | AB,<br>tg(*kop:lifeact.EGFP.nos3'UTR-*<br>*cry:DsRed*) and<br>tg(*kop:GFP.F.nos3'UTR-*<br>*cry:DsRed*) |
| *kop:mCherry.Clip170H.nos3'UTR-*<br>*cmlc:EGFP* (D376) | AB,<br>tg(*kop:lifeact.EGFP.nos3'UTR-*<br>*cry:DsRed*) and<br>tg(*kop:GFP.F.nos3'UTR-*<br>*cry:DsRed*) |
| *kop:mTFP1.Septin9a.nos3'UTR-*<br>*cmlc:EGFP* (D453) | AB |

### Embryo microinjections

One-cell stage zebrafish embryos were microinjected into the yolk with 1 nanoliter of either sense mRNA and or a translation blocking morpholino antisense oligonucleotide (MO; GeneTools). Messenger RNA was synthetized using the mMessageMachine kit (Ambion). The experiment and control embryos were derived from same clutch of eggs. A list of all the constructs used in this study and the mRNA concentration used are provided in Supplementary Table 2 and 3. To express proteins preferentially in germ cells the corresponding coding regions were cloned upstream of the 3’UTR of *nanos3*^49^. For global protein expression the coding regions were cloned upstream of a 3’UTR of the *Xenopus globin* gene. Embryos were kept in 0.3x Danieau’s solution [17.4mM NaCl, 0.21mM KCl, 0.12mM MgSO_4_·7H_2_O, 0.18mM Ca(NO_3_)_2_, 1.5mM HEPES (pH 7.6)] and incubated at 25 degrees prior to imaging.

### Image acquisition

Fluorescence microscopy was performed using a Carl Zeiss Axio imager Z1 equipped with a Retiga R6 camera or a Carl Zeiss Axio imager Z1 equipped with a Spot Pursuit 23.0 1.4 MP monochrome camera. Spinning disk microscopy was performed using a Carl Zeis Axio imager Z1 coupled to a Yokogawa CSUX1FW-06P-01 spinning disk unit and a Hamamatsu Orca-Flash4.0LT C11440 camera. Time lapse movies were acquired using the VisiView software. The confocal microscopy imaging was performed using a Carl Zeiss LSM 710 microscope and the time lapse movies were generated using the Zen software. Imaging of migrating PGCs was performed between shield stage and 90% epiboly. Frames were captured at 8 seconds intervals for experiments performed without the chemokine and 10 seconds intervals were used for the experiments with the chemokine source, with the exception of experiments in Fig. 2a and 2c in which frames were taken every 2 and 7.75 seconds respectively. Embryos were kept at 28 degrees during image acquisition employing a Pecon TempController 2000-2.

### Quantification of bleb location and size

The location of blebs was analyzed by determining the position of the bleb center relative to the cell center. The center of mass of the cell was measured using the FIJI software by drawing the shape of the cell. The center of cell front was defined based on the center of mass of the actin brushes (labeled by Lifeact-GFP). Bleb perimeter was drawn based on the membrane signal (labeled with farnesylated mCherry) and the center of mass was measured in FIJI. The angle between the line connecting the center of mass of the bleb to the center of mass of the cell and the line connecting the center of the mass of the front to the center of mass of the cell was measured in FIJI. Front blebs were defined as having angles smaller than 45 degrees, and rear blebs as showing angles larger than 135 degrees. Angles between 45 and 135 degrees were considered side blebs. To measure the size of the blebs, the bleb perimeter was drawn based on the membrane signal and the area was measured in FIJI.

### Imaging of self-organizing polarity in PGCs

PGCs were imaged between shield and 75% epiboly using widefield or spinning disk microscopy. One-cell stage AB embryos were injected with RNA coding for the specific polarity markers and translation blocking morpholino antisense oligonucleotides directed against Cxcl12a^20^ (GeneTools) (Supplementary Table 4). Images were processed using FIJI software (National Institute of Health). For quantification of the polarization times, time zero was considered to be when the appearance of the actin brushes was first detected. The cascade of appearance of blebs and polarization of other proteins and structures (Ezrin, Esyt2a, Septin9a and MTOC) was followed. At 24 hours post fertilization (hpf), injected embryos were checked for ectopic PGCs, to confirm that the morpholino directed against Cxcl12a functioned well.

### Imaging of chemokine-dependent polarity in PGCs

These experiments were performed by transplantation of zebrafish somatic cells expressing Cxcl12a into host embryos depleted of this chemokine. PGCs of host embryos were either transgenic or injected with RNA coding for the different polarity markers. The host embryos were injected with morpholino against Cxcl12a to deplete the endogenous chemokine and ensure migration towards the transplanted Cxcl12a-expressing cells. Donor embryos were injected two hours after the host embryos with RNA leading to global expression of the chemokine Cxcl12a and of a marker for somatic nuclei or membrane. Cells from donor embryo (between 30% epiboly and shield stages) were transplanted into host embryo (between shield and 75% epiboly) on a Carl Zeiss Axio imager Z1 microscope with a 5x objective. The transplantation was immediately followed by the acquisition of a time lapse video using either 40x or a 63x objective. Image processing and quantification was performed as described above for the self-organizing polarity. At 24 hpf, host and donor embryos were checked for ectopic PGCs to confirm that the morpholino directed against Cxcl12a and the global Cxcl12a expression indeed resulted in ectopic distribution of the cells.

### Photoactivatable Rac1 experiments

Morpholino antisense oligonucleotide directed against Cxcl12a, RNA encoding for *Ezrin-YPet.nos3’UTR* or *Ypet.esyt2al.nos* and either a photoactivatable version of Rac1 fused to the 3’UTR of *nanos3* or control RNA, were injected in one-cell stage *kop:lifeact.mCherry.nos3’UTR-cry:DsRed* embryos. 63X movies of PGCs were acquired using a Carl Zeiss LSM 710 confocal microscope and the photoactivation was performed using the bleaching module in the Zen software. Photoactivation was started after 5 frames with the 456 nm laser using a circular ROI with a diameter of 5 *µ*m (120-240 iterations, 100% laser power) and was repeated every 4 frames. The ROI was located at the edge of the cell opposite to existing actin brushes or a random location in apolar cells.

### DN-Rac1, DN-ROCK, CA-RhoA and hyperactive Ezrin Experiments

One-cell stage embryos of transgenic fish expressing polarity markers or AB fish were injected with RNA encoding for polarity markers and for the specific dominant negative or constitutively active protein fused to the *nanos* 3’ UTR to direct their expression to PGCs. Morpholino directed against *Cxcl12a* was also injected. The constructs and the amount of RNA injected are presented in Supplementary Tables 1 and 2. PGCs were imaged between shield stage and 90% epiboly either in the absence of Cxcl12, or in the presence of transplanted cells expressing the chemokine.

### Direction of the front protrusion of PGCs expressing CA-RhoA relative to the chemokine gradient

The direction of the front protrusion of PGCs expressing CA-RhoA was analyzed by measuring the angle between the line connecting the position of the chemokine source to the center of the cell and the line connecting the center of the cell to the stable protrusion in the last frame of each time lapse movie. An angle of 0**°** indicated perfect alignment of the protrusion towards the chemokine source. The angles were plotted into a rose plot generated in Python.

### Deconvolution

The time lapse movies from Figures 1f, 1g, 2a and Supplementary Figure 2 were deconvolved using software Huygens professional.

### Statistics and reproducibility

Statistical analysis was conducted using Graphpad Prism 6.0d software (Inc., La Jolla, CA). The statistical test used and the *n* number with the p-values are indicated in the figure legends. All experiments were repeated at least three times.

## Acknowledgements

This work was supported by the European Research Council (ERC, CellMig, no. 268806), the Deutsche Forschungsgemeinschaft (DFG, RA863/11-1 and SFB 1348), and the Cells in Motion Cluster of Excellence (EXC 1003-CIM). We thank Nina Knübel for the initial design of the model and Celeste Brennecka for critical reading of the manuscript. We thank Dr. Bart Vos for the Python script for the rose plot and Ina Halbig, Ursula Jordan, Esther-Maria Messerschmidt and Ines Sandbote for technical help.

## Author Contributions

A.O. performed the experiments and analyzed the data shown in Fig. 1b,c, f-p; Fig. 2, Fig. 3 and Supplementary Fig. 2 and cloned DNA constructs. A.A. performed the experiments and analyzed the data shown in Fig. 1a,d and e; Fig. 4 and Fig. 5, Supplementary Fig. 1 and Supplementary Fig 3 and cloned DNA constructs. B.M performed preliminary experiments, performed the experiment in Fig. 2a and cloned DNA constructs. M.R.-F. Cloned DNA constructs. A.O., A.A., designed the experiments with help from B.M., M.R-F. and E.R.. A.O., A.A., and E.R. wrote the manuscript. E.R. supervised the project. All authors read, commented and approved the manuscript.

## Competing Interests statement

Authors have no conflict of interest.

---

**Supplemental Fig. 1.**
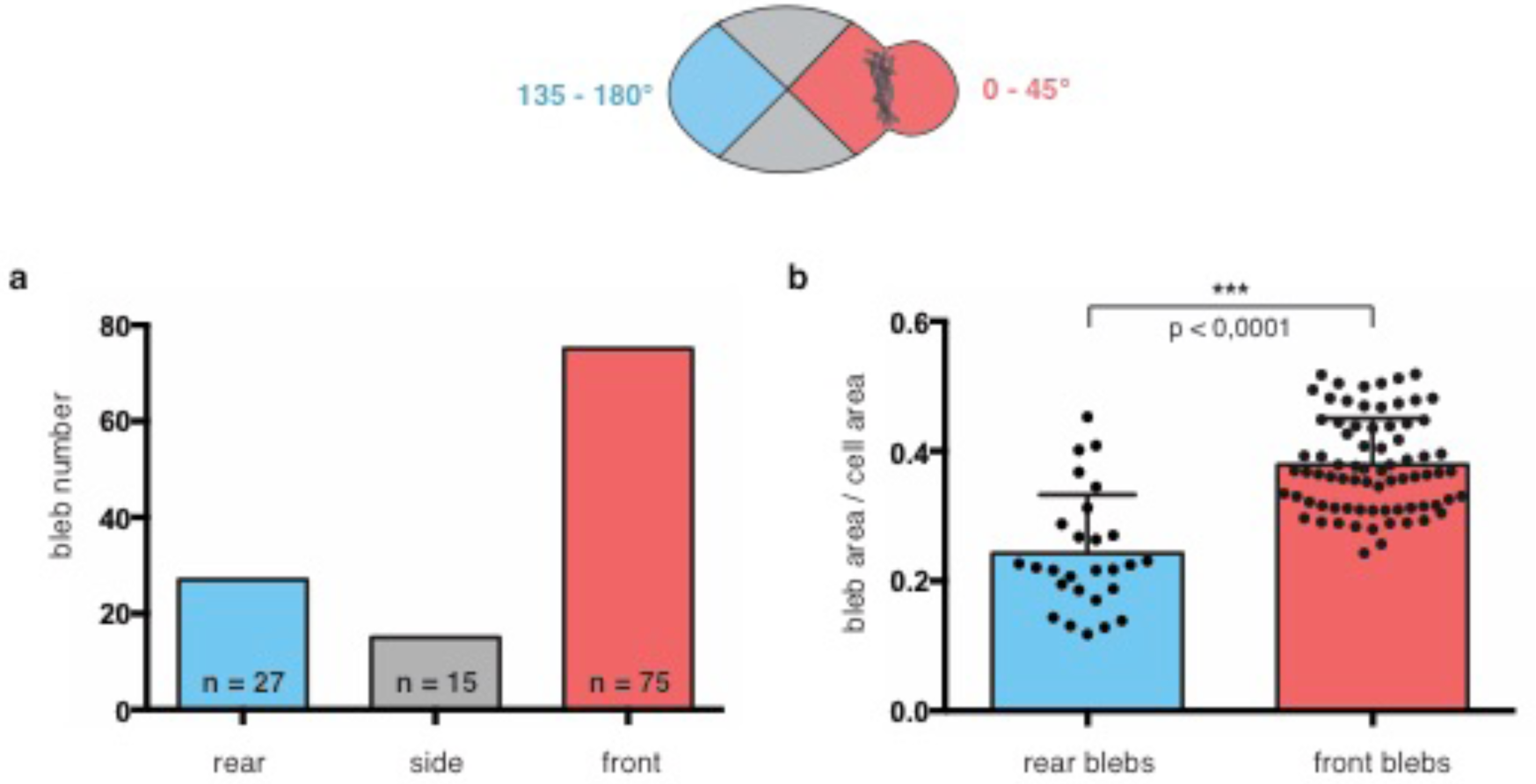
The distribution and size of blebs in wild type migrating PGCs. **a** The number of blebs in a polarized wild type PGCs (n = number of blebs, data obtained from 132 blebs in 19 cells). **b** Front blebs (red) are larger in size relative to the size of the cell as compared with back blebs (blue) (two-tailed t-test, p,< 0.0001).

**Supplemental Fig. 2.**
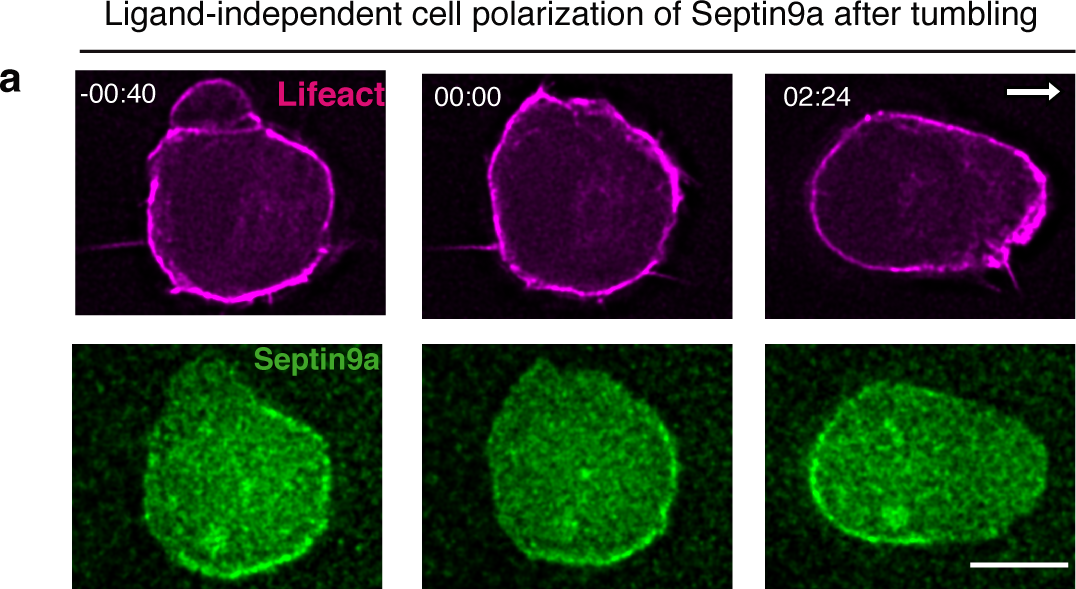
The dynamics of Septin9a localization during self-organizing cell polarization in the absence of receptor signaling. **a** Left panels show an apolar cell, middle panels show the appearance of actin brushes (magenta) in the new front (considered as time “00:00”) and the right panels show the polarized cell with the Septin9a protein (green) localized to the cell rear. White arrow indicates the direction of migration. Time in minutes and seconds. Scale bar 10 *µ*m.

**Supplemental Fig. 3.**
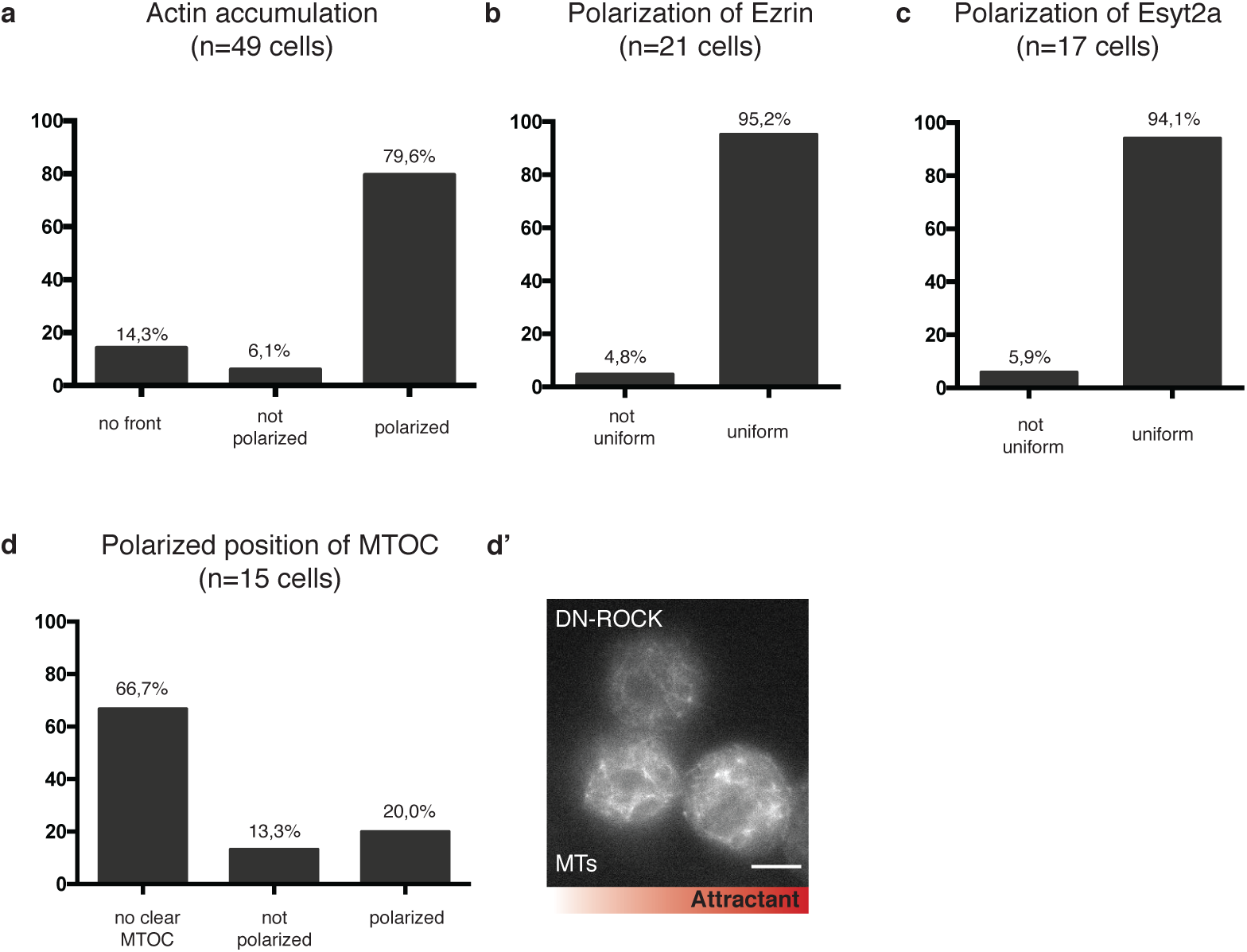
Inhibition of contractility prevents the polarization of the rear markers in the presence of the chemokine. **a-d** Chemokine-directed cell polarization in the presence of DN-ROCK. **a** Formation of a cell front (localized Lifeact signal in the direction of the chemokine source) is not affected by inhibition of contractility. Lack of contractility affects the polarization of **b** Ezrin, **c** Esyt2a and **d** MTOC. **d’** A representative image of germ cells expressing the DN-ROCK with the MTs labeled with a Clip170 fusion protein. The source of the chemokine is on the right side of the panel as illustrated by the red gradient. Scale bar 10*µ*m.

